# Development of a versatile LCM-Seq method for spatial transcriptomics of fluorescently-tagged cholinergic neuron populations

**DOI:** 10.1101/2023.03.02.530916

**Authors:** Éva Rumpler, Balázs Göcz, Katalin Skrapits, Miklós Sárvári, Szabolcs Takács, Imre Farkas, Szilárd Póliska, Márton Papp, Norbert Solymosi, Erik Hrabovszky

## Abstract

Single-cell transcriptomics are powerful tools to define neuronal cell types based on co-expressed gene clusters. Limited RNA input in these technologies necessarily compromises transcriptome coverage and accuracy of differential expression analysis. We propose that bulk RNA-sequencing of neuronal pools defined by spatial position offers an alternative strategy to overcome these technical limitations. We report an LCM-Seq method which allows deep transcriptome profiling of fluorescently-tagged neuron populations isolated with laser-capture microdissection (LCM) from histological sections of transgenic mice. Mild formaldehyde-fixation of ZsGreen marker protein, LCM sampling of ∼300 pooled neurons, followed by RNA isolation, library preparation and RNA-sequencing with methods optimized for nanogramm amounts of moderately degraded RNA enabled us to detect ∼15,000 different transcripts in fluorescently-labeled cholinergic neuron populations. The versatile LCM-Seq method showed excellent accuracy in quantitative studies, with 2,891 transcripts expressed differentially between the spatially defined and clinically relevant cholinergic neuron populations of the caudate-putamen and medial septum.

## Introduction

Genome-wide quantification of mRNA transcripts is a powerful approach to characterize cellular states and molecular circuitries. High throughput next-generation sequencing of RNA (RNA-Seq) increasingly replaces hybridization-based microarray technologies (1) given its much better dynamic range, higher reproducibility and increased accuracy in differential expression analysis of both high- and low-abundance transcripts (2).

*Spatially resolved transcriptomics* includes a growing variety of RNA-Seq technologies which preserve topographic information on the analyzed cells (3-7). Considering the extraordinary diversity of brain cell types, retaining the precise anatomical context is particularly important in neuroscience (8). Spatial transcriptomic methods are often complementary, with various inherent strengths and trade-offs in anatomical resolution, target coverage and throughput. MERFISH (9) and other *in situ* single-cell transcriptomics (scRNA-Seq) which use microscopic analysis of multiplexed fluorescence *in situ* hybridization (FISH) signals, provide unparalleled anatomical precision, with a necessary compromise in the detection of low-abundance transcripts. A large number of scRNA-Seq techniques use simultaneous *ex situ* sequencing of thousands of dispersed cells (10-13). While this is an efficient approach to reveal neuronal heterogeneity and define cell types (14, 15), the limited amount of RNA in single cells (16) compromises detection sensitivity, accuracy and precision and reduces the statistical power of differential gene expression analysis (10-12).

Bulk sequencing of neuronal pools can overcome several limitations of scRNA-Seq methods by increasing the amount of starting RNA (11, 17, 18). Use of laser capture microdissection (LCM) (19) to collect a spatially defined cell population from frozen tissue sections, coupled to RNA-Seq (LCM-Seq) (17, 20, 21) is a powerful strategy to obtain a transcriptomic snapshot from anatomically similar cell populations. Accordingly, application of morphometric selection criteria for LCM has recently allowed us to carry out deep transcriptome profiling of spiny projection neurons and cholinergic interneurons (ChINs) from the human putamen (17). Adaptation of the LCM-Seq technology to neuronal populations expressing specific neurochemical phenotype markers remains technically challenging and unresolved.

To address an unmet need, here we introduce a versatile LCM-Seq method for deep transcriptome analysis of spatially defined neuron populations expressing a fluorescent marker protein. Use of ZsGreen reporter protein fixed mildly with formaldehyde, LCM sampling of >300 pooled neurons, and RNA isolation, library preparation and RNA-sequencing methods optimized for nanogramm amounts of moderately degraded RNA enabled us to detect ∼15,000 different transcripts in distinct cholinergic neuron populations of transgenic mice. Beyond this high sensitivity, the LCM-Seq method showed excellent accuracy and precision in quantitative studies, allowing us to identify 2,891 genes expressed differentially between the spatially defined cholinergic neuron populations of the caudate-putamen (CPU) and the medial septum (MS), two clinically important cell types in various neurodegenerative disorders.

## Results

### ZsGreen fixed lightly with formaldehyde serves as an optimal fluorescent marker for LCM-based cell harvesting

Fluorescent proteins expressed in transgenic mice are widely used neuronal phenotype markers. Harvesting fluorescent neurons with LCM for downstream RNA applications, such as RT-qPCR, microarray and RNA-Seq, requires stabilization of the fluorescent signals with the cross-linking fixative formaldehyde (18, 22, 23). As the first step toward an optimized LCM-Seq strategy, we compared the fluorescence stability and signal intensity of formaldehyde-fixed ZsGreen and tdTomato proteins expressed selectively in cholinergic systems (**Fig. 1a**). We perfused the vasculature of ChAT-tdTomato and ChAT-ZsGreen transgenic mice (N=3/group) transcardially with either 0.5%, 2% or 4% formaldehyde, followed by 20% sucrose. Forebrain sections were mounted on microscope slides and analyzed with fluorescence microscopy. We found that tdTomato fluorescence was extremely weak using 0.5% formaldehyde, with very little improvement using 2% or 4% fixative. In contrast, all three formaldehyde concentrations preserved the bright ZsGreen signal in cholinergic neurons for subsequent isolation with LCM (**Fig. 1a**).

**Fig. 1:**
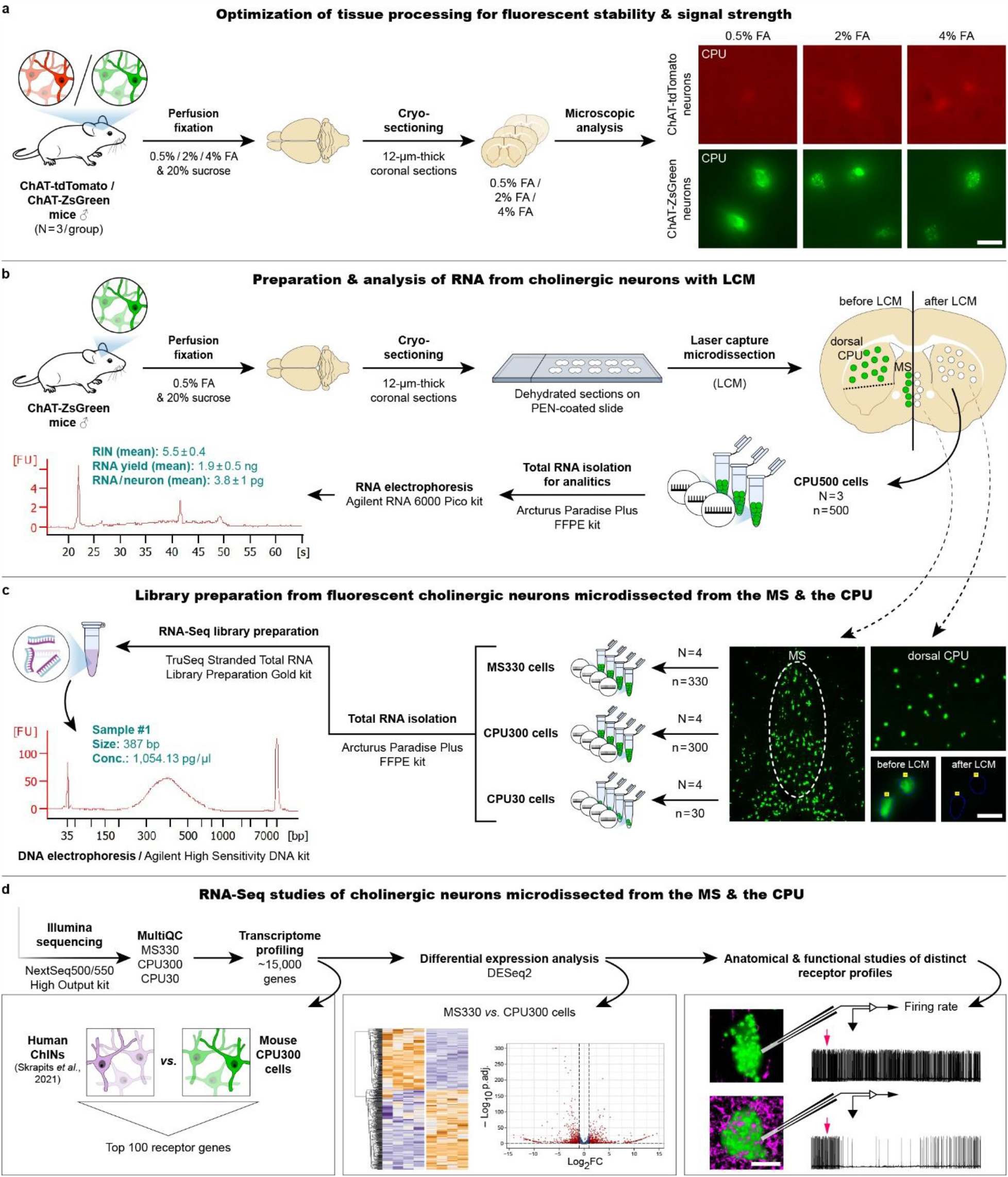
LCM-Seq protocol developed for whole transcriptome profiling of spatially resolved cholinergic neuron populations. **a:** Effect of fixation on marker protein fluorescence. Microscopic analysis of sections obtained from the CPU of ChAT-tdTomato and ChAT-ZsGreen transgenic mice provides evidence for superior signal intensity of formaldehyde (FA)-fixed ZsGreen over tdTomato, irrespective of fixative concentration. **b:** Bioanalyzer measurements on RNA isolated from 500 pooled ChINs. **c:** Random primer-based RNA-Seq library preparation from 330 MS cholinergic neurons (MS330) and 300 CPU ChINs (CPU300) occupying equal section areas. Additional libraries from 30 ChINs (CPU30) are prepared to investigate how limited RNA input affects sequencing quality. **d:** Illumina sequencing and bioinformatic analysis of two spatially distinguished cholinergic neuron populations. Quality control (QC), transcriptome profiling and differential expression analysis are followed by neuroanatomical and functional confirmation of unique receptorial mechanisms revealed in the two distinct cell types. Scale bar: 25 µm (**a** and high-power images in **c**), 200 µm (low-power images in **c**), 10 µm (**d**).

### ChAT-ZsGreen neurons collected with LCM provide ∼4 pg/cell total RNA with a RIN value of ∼5.5

To minimize fixation-related RNA damages (24), we optimized further the LCM-Seq protocol using ChAT-ZsGreen brains fixed lightly with 0.5% formaldehyde. Twelve-µm-thick coronal sections of three mice were mounted on PEN membrane slides. Fluorescent ChINs were microdissected individually and collected from the dorsal CPU with LCM (**Fig. 1b**). To estimate the amount and quality of total RNA recovered from LCM-isolated neurons, we prepared RNA samples (N=3) from 500 pooled neurons with the Arcturus Paradise PLUS FFPE RNA Isolation Kit (ThermoFisher). RNA yield we determined with Bioanalyzer analysis was 1.9±0.5 ng (∼3.8±1 pg/neuron) and RNA integrity number (RIN) used as a sample quality measure was 5.5±0.4 (mean±SEM) (**Fig. 1b**).

### Preparation of RNA-Seq libraries from LCM-isolated neurons combines rRNA removal and random primer-based reverse transcription

Orthodox RNA-Seq library preparation protocols rely on the reverse transcription of mRNA with oligo(dT) primers which becomes inefficient using low-quality and low-quantity input RNA (25). Available commercial kits optimized for low amounts and fragmented RNA require inventive hybridization-based removal of ribosomal RNA (rRNA) or cDNA sequences with highly efficient random primer-based reverse transcription of RNA transcripts (25, 26). From a variety of approaches available (25), we chose to prepare cholinergic cDNA libraries with the TruSeq Stranded Total RNA Library Preparation Gold kit (Illumina, San Diego, CA, USA) which has recently enabled us to bulk sequence 300-400 formaldehyde-fixed neuronal cells from both mice (18) and humans (17) (**Fig. 1c**). To exceed safely the cDNA sample requirements of Illumina sequencing, we increased the number of amplification cycles from 15 to 16 while preparing libraries from 330 LCM-isolated basal forebrain cholinergic cells of the MS (MS330; N=4) and 300 microdissected ChINs of the dorsal CPU (CPU300; N=4). Choosing these cell numbers ensured that MS and CPU neurons occupy identical section areas (CPU300: 66,476±1,796 µm^2^; MS330: 67,666±1,569 µm^2^). To also address the impact of limited RNA input on sequencing quality, we generated and studied additional libraries (CPU30; N=4) from 30 microdissected CPU ChINs, using 19 PCR cycles for fragment enrichment (**Fig. 1c**). The amount of cDNA in the 12 amplified samples varied between 278-5,933 pg/µl. Twenty µl of a 1 nM library mix representing proportionally the indexed samples was subjected to single-end sequencing with the Illumina NextSeq500/550 High Output kit (v2.5; 75 Cycles) and an Illumina NextSeq500 instrument (**Fig. 1d**).

### LCM-Seq studies of 300-330 microdissected neurons allow the identification of ∼15,000 different transcripts in two spatially defined cholinergic neuron populations

Sequencing of the twelve RNA-Seq libraries generated 516.5 M raw reads (27.3 M-48.6 M per sample). After trimming with Trimmomatic and adapter removal with Cutadapt, mapping to the mouse reference genome (Ensembl; release 107) with STAR (v 2.7.9a) resulted in 26±4.7M and 31.5±2.8M aligned reads in the MS330 and CPU300 samples (**Fig. 2a**), respectively, from which 10.3±1.8M MS330 reads and 13.2±2.1M CPU300 reads were assigned to unique genes (**Fig. 2a**), and quantified with FeatureCounts (subread v 2.0.2). For gene body coverage and read distribution between exons, introns and intergenic regions, see **Supplementary Figure 1**. LCM-Seq studies of the 300-330 pooled cholinergic neurons allowed us to detect 14,926±422 different transcripts in MS330 and 15,353±231 transcripts in CPU300 samples (cutoff: 5 reads/sample; **Fig. 2b**). All transcripts identified in MS and CPU cholinergic neurons and selected functional categories, such as *ion channels, transporters, transcription factors* and *receptors* (defined in the KEGG BRITE database) are reported in **Supplementary Table 1**.

**Fig. 2:**
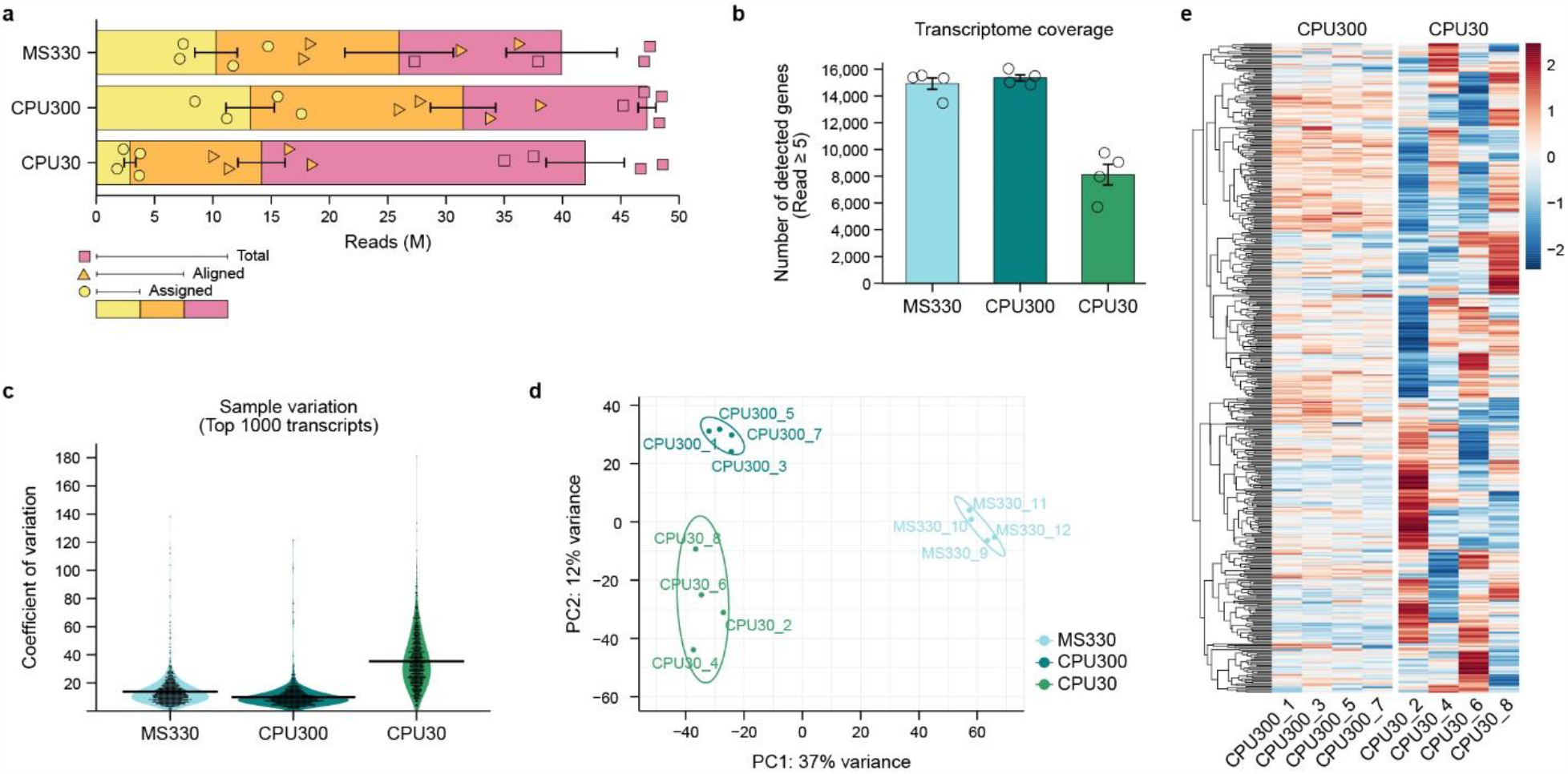
Quality assessment of LCM-Seq results. **a:** Number of total reads, reads alignable to the mouse reference genome (Ensembl; release 107) and reads assigned to unique genes. While total reads are distributed evenly among the co-sequenced 12 samples, there is a large drop in both aligned and assigned reads using RNA harvested from 30 (CPU30 samples), instead of 300-330 pooled cholinergic neurons (CPU300 and MS330 samples; N=4 for each). **b:** Effect of RNA input on library complexity. The 2.9-13.2 M assigned reads allow the identification of ∼15,000 different transcripts (raw reads≥ 5) in the MS330 and CPU300 transcriptomes. For full listing and categorization of these transcripts, see **Supplementary Table 1**. The number of genes detected with the same cutoff is compromised substantially in the CPU30 (N=4) transcriptomes (∼8,100 transcripts). **c:** Coefficient of variation for the most highly expressed 1,000 transcripts in each library. Violin plot illustrates enhanced sample variability in the CPU30 (mean CV: 35.4) *vs*. the MS330 (mean CV: 13.7) and the CPU300 (mean CV: 9.83) transcriptomes. **d:** Principal component plot. Cell type-dependent differences (MS *vs*. CPU neurons) are best reflected in clear separation of the MS330 from both the CPU300 and the CPU30 transcriptomes in PC1. Difference between CPU30 and CPU300 in PC2 reveals a library amplification bias due to low amounts of starting RNA in the former. Note also the reduced clustering of the CPU30 transcriptomes. **e:** Heat map of the top 1,000 transcripts identified in the CPU300 (N=4) and CPU30 (N=4) transcriptomes. Enhanced CPU30 sample variability indicates largely reduced detection accuracy and precision.

### Limited RNA input compromises sequencing quality, library complexity and detection accuracy

While bulk sequencing of 300-330 microdissected neurons ensured excellent transcriptome coverage, we found that several qualitative parameters of sequencing were compromised when the number of pooled neurons was reduced to 30. Accordingly, the number of reads aligned to the mouse genome (**Fig. 2a**), assigned to unique genes (**Fig. 2a**) and mapped to exonic seqences (**Supplementary Figure 1b)** dropped in the CPU30 transcriptomes. The number of different transcripts (cutoff: raw reads ≥ 5) decreased from 15,353±231 in the CPU300 to 8,118.25±768 in the CPU30 transcriptomes (**Fig. 2b**), indicating reduced library complexity. Limited RNA input was also harmful for detection accuracy. The coefficient of variation increased from 9.83 (mean of the most abundant 1,000 transcripts) in the CPU300 to 35.4 in the CPU30 transcriptomes (**Fig. 2c**). Further, CPU30 transcriptomes exhibited impaired clustering in the PCA plot and differed from the CPU300 transcriptomes in PC2 (**Fig. 2d**). Along with an increased color variability of heat maps (**Fig. 2e**), these observations indicated that the low RNA input introduced amplification biases and unwanted variations in the CPU30 libraries.

### Genes implicated in Alzheimer’s disease and Parkinsons’s disease are highly enriched in cholinergic neurons

Cholinergic systems are susceptible to degeneration in Alzheimer’s disease (AD) and Parkinsons’s disease (PD). Analysis of Human Disease Ontology databases revealed an interesting enrichment of AD and PD genes in the MS330 and CPU300 transcriptomes. Half of the genes listed in AD (N=156) and PD (N=133) databases were present (>50 CPM) and ∼27-35% were highly expressed (>50 CPM) in the MS330 and CPU300 transcriptomes (**Supplementary Figure 2)**.

### Human and murine striatal ChINs exhibit conserved receptor profiles

We have reported recently the detailed gene expression profile of LCM-isolated human ChINs identified by size in cresyl violet stained putamen sections (17). This allowed us to address similarities/differences between the functionally analogous striatal neurons of the two species. The most abundant 100 receptor transcripts (identified based on the KEGG BRITE database), including the *Drd2* and *Drd5* dopaminergic receptor isoforms, were expressed at strikingly similar levels in humans and mice (**Fig. 3**). This finding revealed evolutionarily conserved regulatory mechanisms acting in striatal ChINs. The few exceptions showing species-specific expression included *Drd3* in human and the opioid receptor *Oprl* in murine ChINs (**Fig. 3; Source Data 1**).

**Fig. 3:**
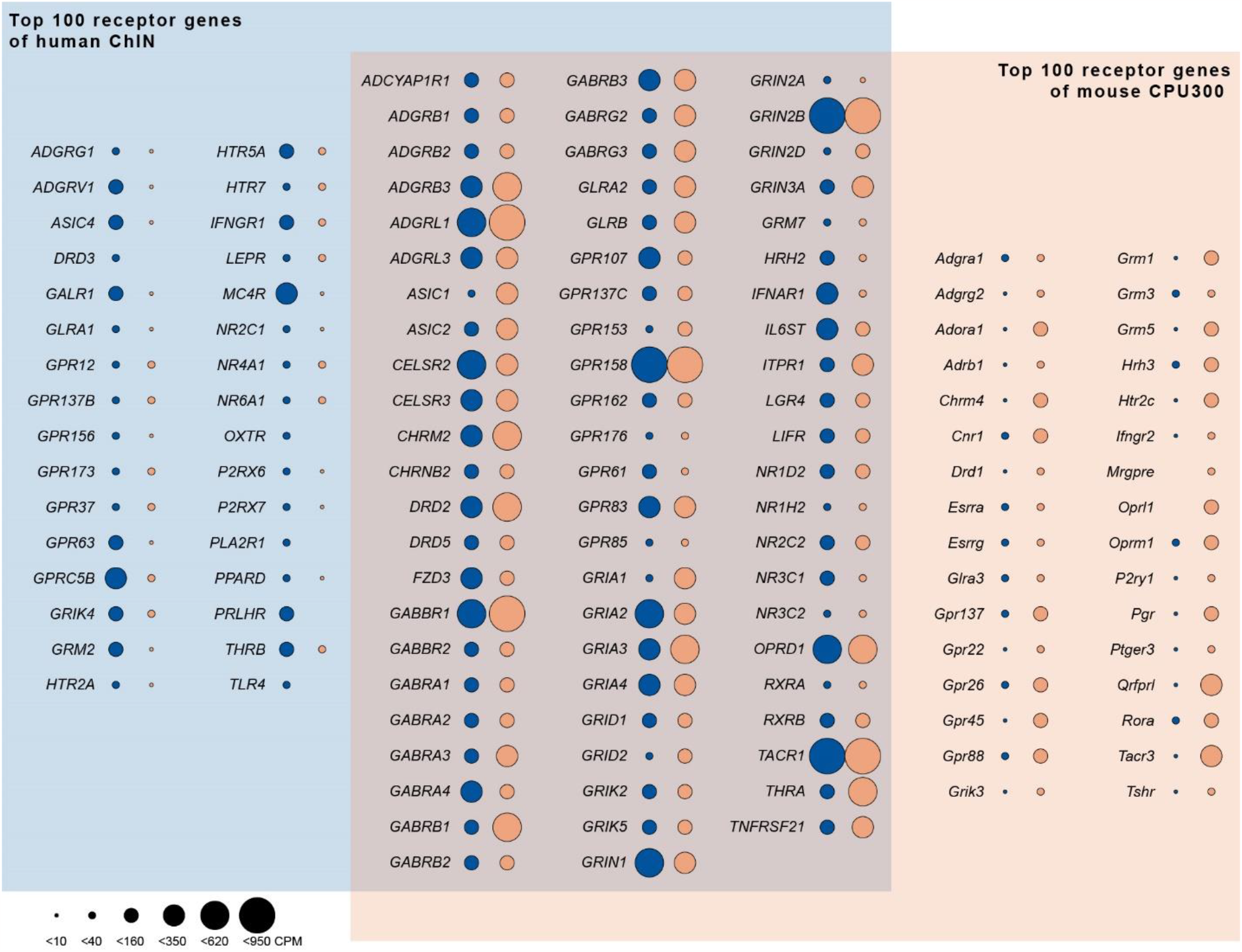
Receptor expression landscape of striatal ChINs from the human putamen and the mouse CPU. ChINs of the human putamen (blue field) (17) and the murine CPU (orange field) exhibit striking similarities in receptor expression profiles. Venn diagram illustrates the most abundant 100 receptors (KEGG BRITE database) from each species. Dark blue (human) and dark orange (mouse) dot areas change in proportion to transcript abundances [counts per million (CPM) units]. Items in the overlapping area occur among the top 100 ChIN receptors in both species. Note obvious species similarities in all three fields and the few receptors detected only in human or murine ChINs. See also **Source Data 1**.

### 2,891 transcripts are expressed differentially in the MS330 and CPU300 transcriptomes

Both the MS330 and the CPU300 transcriptomes contained cholinergic as well as GABAergic neuronal phenotype markers (**Fig. 4a**), indicating simultaneous use of acetylcholine and GABA for neurotransmission. Differential expression analysis of the two transcriptomes using the DESeq2 R-package with RUVSeq normalization revealed 2,891 transcripts with significantly different levels (p.adj.<0.05) between MS and CPU neurons (**Fig. 4b; Source Data 2**). Homogenous heat maps of these transcripts (**Fig. 4c**) indicated excellent accuracy and precision of the LCM-Seq approach for studies of differential gene expression.

**Fig. 4:**
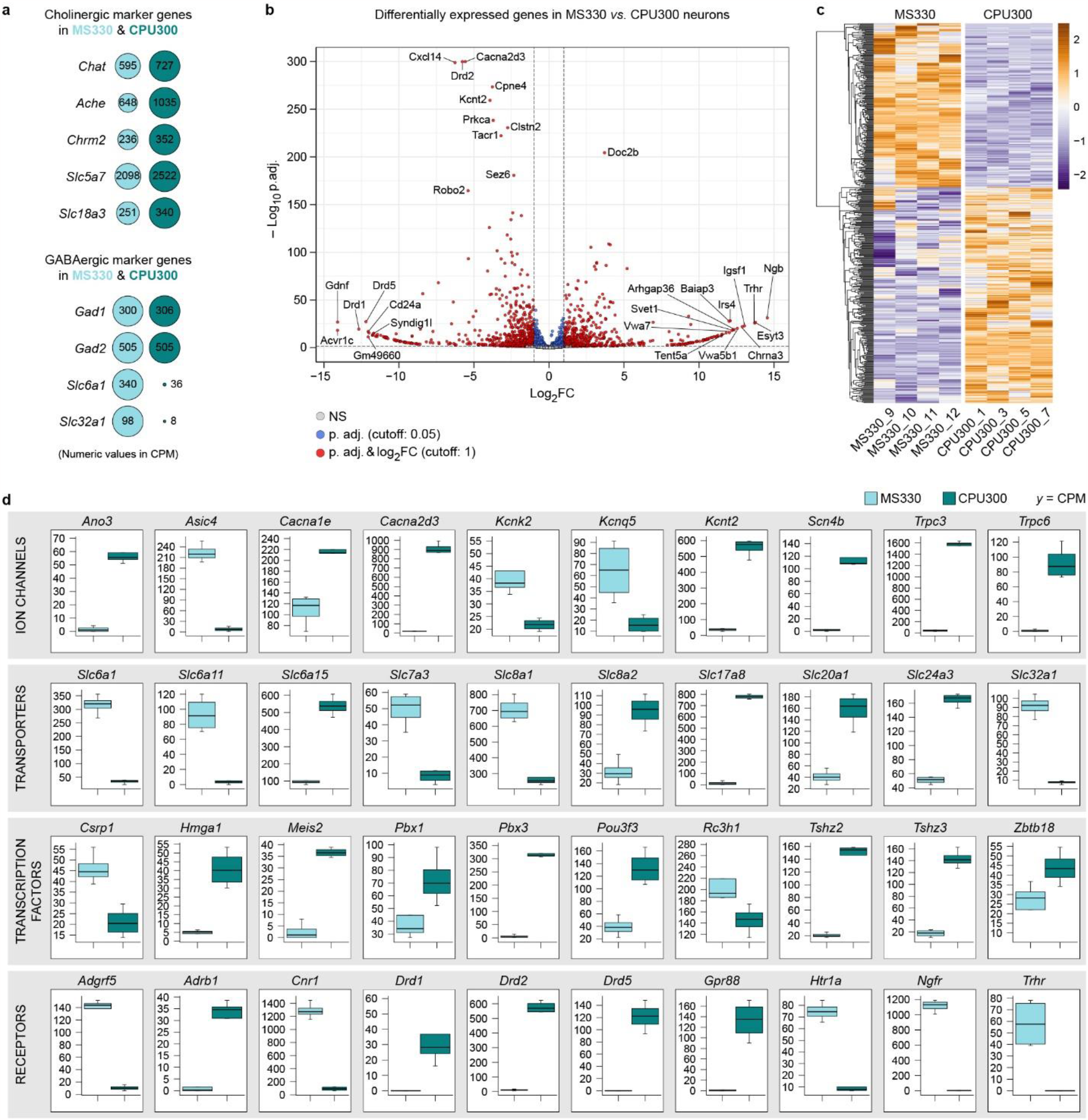
Differential gene expression analysis of cell type specific regulatory mechanisms acting in MS and CPU cholinergic neurons. **a:** Expression of known cholinergic and GABAergic phenotype markers in each cell type. **b:** Differentially expressed transcripts illustrated in a volcano plot. Genes with |log2 fold change (FC)|>12 and -log_10_ p.adj>150 are tagged with gene symbols. **c:** Heat map of 2,891 transcripts expressed at significantly different levels in the MS330 and CPU300 transcriptomes (p.adj<0.05). **d:** Tukey’s box plots with examples of differentially expressed ion channels, transporters, transcription factors and receptors. For full listing, see **Source Data 2**. Abundant expression of *Drd2* only in CPU and *Trhr1* only in MS cholinergic neurons forecast differential responsiveness of the two cell types to D2 agonists and to TRH, respectively.

### MS and CPU cholinergic neurons have unique ion channel, transporter, transcription factor and receptor profiles

Cholinergic projection neurons of the MS and the diagonal band of Broca give rise to the septohippocampal pathway. This connection plays a critical role in hippocampal theta rhythms and memory encoding, among other functions (27, 28). Cholinergic neurons of the CPU, in turn, contribute as local interneurons to the regulation of cortico-striato-thalamocortical neural pathways and are involved in motor control, learning, language, reward, cognitive functioning, and addiction (29). Different regulatory functions of MS and CPU neurons were reflected in 108 ion channels, 111 transporters, 140 transcription factors, and 81 receptors among the 2,891 transcripts we found to be expressed differentially (p.adj.<0.05) between the MS330 and CPU300 transcriptomes (**Fig. 4d; Source Data 2**). Using Synaptic Gene Ontology (SynGO) pathway analysis, we identified hundreds of differentally expressed transcripts related to various aspects of synaptic neurotransmission (**Supplementary Figure 3; Source Data 2)**.

### Cell type-specific receptor profiles are in line with the distinct functional properties of MS and CPU cholinergic neurons

Differential expression analysis revealed selective presence of the *Drd1, Drd2* and *Drd5* dopamine receptor isoforms in CPU cholinergic neurons (**Fig. 4d**), suggesting that dopamine plays a key role in the afferent control of this cell group. Indeed, the immunofluorescent detection of the dopaminergic marker enzyme tyrosine hydroxylase (TH) revealed a dense dopaminergic fiber network in the CPU. In contrast, we could only label a few TH fibers in tissue sections of the MS (**Fig. 5a**). Corroborating distinct *Drd2* expression profiles and TH innervation patterns, whole-cell current-clamp studies of ChAT-ZsGreen neurons from the CPU established that a single bolus of the D2 agonist sumanirole (30 μM) decreased the rate of cholinergic neuron firing to 25.7±5.8% (mean ± SEM) of the control. In contrast, the mean firing frequency of MS cholinergic neurons remained unaffected (108.6±7.5% of the control frequency) (**Figs. 5b, c; Source Data 3**), in accord with the extremely low *Drd2* expression in this cell type (**Fig. 4d**).

**Fig. 5:**
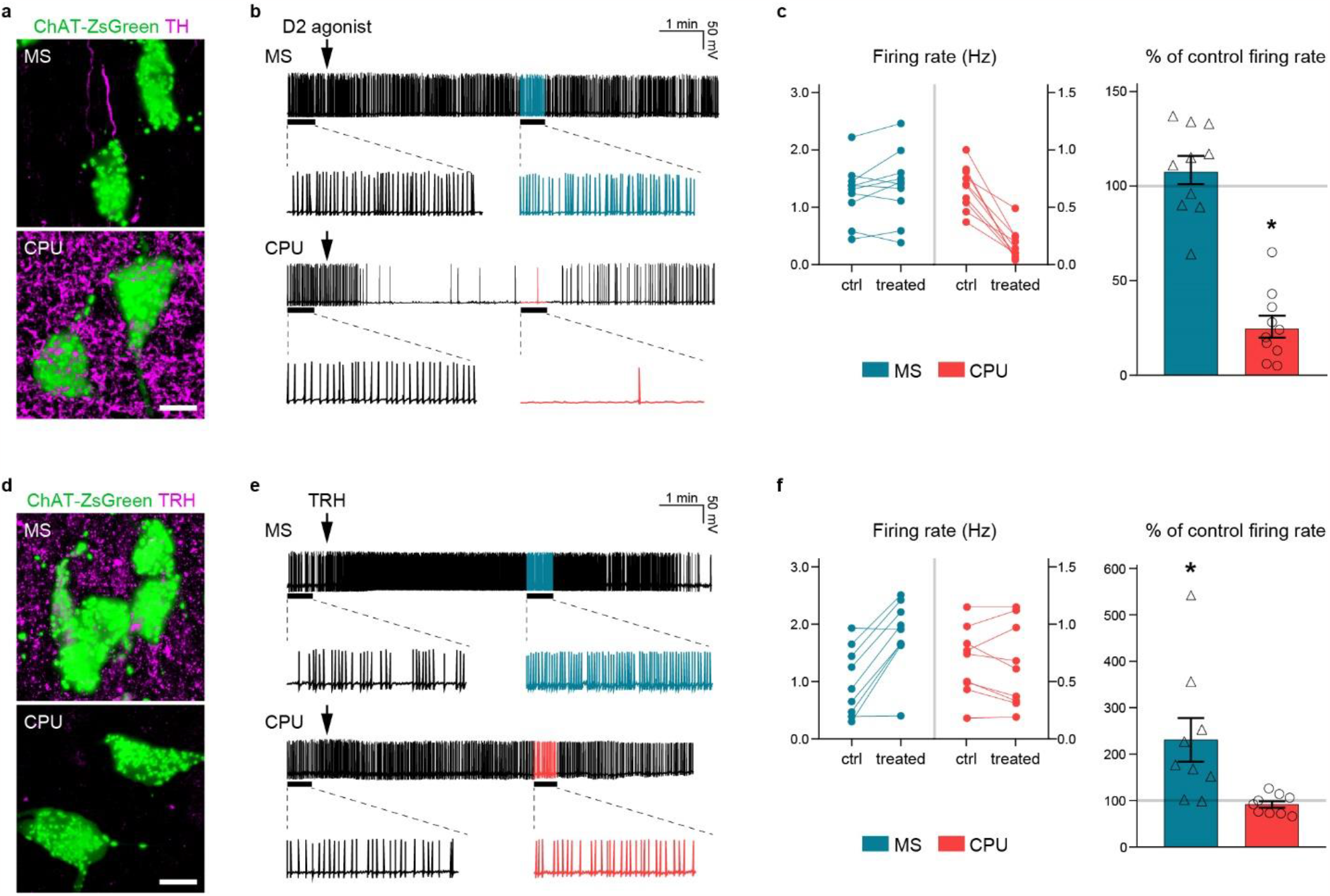
Anatomical and functional support for differential responsiveness of MS and CPU cholinergic cell types to a D2 agonist and to TRH. **a:** Confocal microscopic analysis of ChAT-ZsGreen cholinergic neurons (green) and dopaminergic fibers immunoreactive to tyrosine hydroxylase (TH; magenta). While dopaminergic axons are scarce in the MS, they form a dense network around ChAT-ZsGreen neurons of the CPU, suggesting strong dopaminergic control of the latter cell type. **b, c:** Results of whole cell patch clamp recordings from MS and CPU cholinergic neurons treated with a single bolus of the selective D2 receptor agonist sumanirole (30 μM). While representative traces (**b**) and statistical analyses (**c**) reveal no change in the mean firing frequency of MS cholinergic neurons, sumanirole causes robust inhibition of CPU cholinergic neurons. These observations are in line with high expression of *Drd2* only in the latter cell group. **d:** Simultaneous detection of ChAT-ZsGreen cells (green) and TRH immunoreactive axons (magenta). TRH axons are present in high numbers and surround cholinergic neurons in the MS, whereas they occur rarely in sections of the CPU. **e, f:** Electrophysiological responses of MS and CPU cholinergic neurons to TRH in representative traces (**e**) and in statistics (**f**). A single bolus of TRH causes robust excitation of cholinergic neurons in the MS, without affecting the mean firing rate of CPU cholinergic neurons. This observation is in accord with the selective RNA-Seq detection of *Trhr1* in the former cell group. Scale bars: 10 μm. * p<0.05 *vs*. ctrl with Student’s t-test. For statistics, see **Source Data 3**.

Among the differentially expressed receptors, the TRH receptor *Trhr1* showed overwhelming expression in the MS330 transcriptome (**Fig. 4d**), suggesting differential TRH responsiveness of the two cholinergic cell types. As predicted, neuronal fibers immunoreactive to proTRH occurred in high numbers in the MS (**Fig. 5d**). Interestingly, a subset (8.4±2.1%) of MS cholinergic neurons showed immunoreactivity to proTRH (**Supplementary Fig. 4a, b**). Indeed, *Trh* mRNA expression in the MS330 transcriptome could be verified (**Supplementary Fig. 4c** and **Supplementary Table 1**). Identification of endogenous TRH and TRHR1 in a subset of MS cholinergic neurons raised the intriguing possibility of an autocrine regulatory mechanism acting in these cells. Cholinergic neurons in the CPU, unlike in the MS, were not surrounded by TRH immunoreactive fibers (**Fig. 5d**). Using slice electrophysiology, we established that a single bolus of TRH (10 μM) increased the firing rate of MS cholinergic neurons to 230.7±47.2% of the control rate. In contrast, the mean firing rate of CPU cholinergic neurons, lacking *Trhr1* expression, remained unaffected (91.7±7.1% of the control rate) (**Figs. 5e, f; Source Data 3**). Altogether, these immunohistochemical and electrophysiological observations provided functional support for different regulatory mechanisms acting in the two spatially defined cholinergic cell types.

## Discussion

In this study we are reporting a versatile LCM-Seq method developed for deep transcriptome profiling of fluorescently-tagged neuron populations isolated with laser-capture microdissection (LCM) from histological sections of transgenic mice. Key findings are that, i) Light formaldehyde fixation preserves the bright fluorescence of expressed ZsGreen, but not tdTomato, during tissue preparation; ii) ChAT-ZsGreen neurons isolated with LCM from 12-µm-thick sections of the dorsal CPU provide ∼4 pg/cell total RNA with only minimally compromised integrity (RIN: ∼5.5); iii) Bulk RNA-sequencing of RiboZero libraries prepared from 300-330 LCM-isolated cholinergic neurons (∼1.2 ng total RNA) can reliably identify ∼15,000 different transcripts with sufficient accuracy and precision to allow differential expression analysis of the two spatially defined cholinergic cell types of the MS and the CPU; and finally, iv) Both detection sensitivity and accuracy of RNA-Seq become compromised using only 30 neurons to prepare the libraries. This versatile LCM-Seq method has unique advantages over various scRNA-Seq technologies (10-13) due to the use of ng, instead of pg (16) amounts of RNA, which can substantially improve detection sensitivity, accuracy and precision (11). Conversely, we have shown formally that the number of the identified transcripts drops from ∼15,000 to ∼8,100, the coefficient of variation for the top 1,000 transcripts increases and PCR amplification of the cDNA libraries introduces biases to the LCM-Seq results if RNA samples are prepared from 30, instead of 300-330 pooled neuronal cells.

Fluorescence activated cell sorting (FACS) of dispersed living neurons (30) offers an alternative protocol to LCM for cell enrichment. One major advantage of LCM over FACS is the superior anatomical precision of sample collection. In addition, the LCM-Seq approach provides a functional snapshot on the transcriptome, whereas the lengthy FACS protocol may allow unwanted transcriptomic changes to occur. Stabilizing the transcriptome with our approach will be critically important in future studies of differential gene expression. Accordingly, the LCM-Seq method has recently allowed us to identify ∼2,300 significant changes in the kisspeptin neuron transcriptome of gonadectomized female mice in response to estrogen treatment (18).

Alcohol-based precipitating (molecular) fixatives were unable to preserve the signals of expressed fluorescent proteins in our pilot studies and the use of formaldehyde in the LCM-Seq protocol was inevitable. Although this crosslinking fixative was reported to reduce RNA quality and recovery (31) and to interfere with the RT-PCR amplification of RNA extracts (24), the amount (∼1.2 ng) and integrity (RIN∼5.5) of RNA we were able to isolate from ∼300 microdissected neurons satisfied requirements of library preparation protocols optimized for low-quality/low-quantity RNA samples (25, 26). The TruSeq Stranded Total RNA Library Preparation Gold kit we chose here relies on the RiboZero technology to remove rRNA, followed by reverse transcription of RNA with random primers and cDNA fragment enrichment with PCR. This procedure requires >100 ng RNA for optimal results according to the manufacturer (Illumina). However, it was tested and reported to perform very well using 5-10 ng, and well using 1-2 ng degraded RNA (26). Our decision to pool ∼300 neurons for the LCM-Seq method relies on practical considerations: i) ∼300 pooled CPU neurons were enough to provide >1 ng total RNA, the minimal amount tested successfully by other investigators (26); ii) Collecting thousands of neurons within a workday with LCM would be technically infeasible; iii) Preparation of libraries from 30 neurons greatly reduced detection sensitivity and precision; iv) Genetically tagged neuron populations with a strictly defined topography often consist of a few hundred microdissectable cells only (18, 23).

In this study we used the new LCM-Seq method to characterize the transcriptome landscape of two clinically important cholinergic neuron populations in the MS and the CPU, respectively. Information we are reporting on ∼15,000 expressed genes in each cell type contribute to our understanding of the molecular regulation of the two distinct neuronal systems in normal and disease conditions. Combined results of RNA-Seq, immunofluorescence and electrophysiological studies supported the critical involvement of dopamine and specifically, the most abundant dopaminergic receptor, D2 in the inhibitory control of striatal ChINs.

Future LCM-Seq studies of ChINs will be able to address the molecular consequences of losing this dopaminergic regulation in animal models of Parkinson’s disease.

Important functions associated with cholinergic neurons of the MS and other basal forebrain regions include cortical activation, affect, attention, sensory coding, motivation, and memory and the demise of this system has been strongly implicated in the pathophysiology of Alzheimer’s disease (28). Basal forebrain cholinergic neurons consist of heterogeneous subpopulations which differ in cortical, amygdalar and hippocampal projections as well as in sources and neurochemistry of their synaptic afferentation (28). LCM-Seq studies of subpopulations will require the identification and selective transgenic expression of marker genes for distinct cell types. In this study we carried out transcriptome profiling of ChAT-ZsGreen neurons from the MS. Out of ∼15,000 expressed transcripts, we selected *Trhr1* for anatomical and electrophysiological studies and provided evidence for a strong excitatory regulation of MS cholinergic neurons by TRH. The unexpected finding of TRH co-synthesis in a subset of cholinergic neurons indicates that some of the receptor ligand may originate from endogenous sources. TRH agonists emerged as promising therapeutics to restore cholinergic functions in various neurodegenerative disorders (32, 33). The well-established analeptic effects of TRH was attributed, at least partly, to TRH activation of the septohippocampal cholinergic pathway over 40 years ago (34). Our study now provides solid molecular, anatomical and functional support to the role of TRH signaling in the excitatory regulation of the septohippocampal cholinergic pathway.

In conclusion, this versatile LCM-Seq method allowed deep transcriptome profiling of fluorescently-tagged neuron populations isolated with LCM from histological sections of transgenic mice. Bulk sequencing of neuronal pools defined by identical spatial position showed excellent sensitivity as well as accuracy and precision in quantitative studies on the clinically relevant cholinergic neuron populations isolated from the MS and the dorsal CPU regions.

## Materials and Methods

### Animals

Genetically modified prepubertal (PND 20-30) and young adult (PND 60-90) male mice (N=31) were housed under standard conditions (lights on between 0600 and 1800 h, temperature 22±1 °C, chow and water *ad libitum*) in the animal facility of the Institute of Experimental Medicine. ChAT-tdTomato and ChAT-ZsGreen transgenic mice were generated by crossing male ChAT-IRES-Cre knock-in mice (Jackson Laboratory, Bar Harbor, ME; RRID: IMSR_JAX:006410) with female B6.Cg-Gt(ROSA)26Sortm14(CAG-tdTomato)Hze/J (Jackson Laboratory, IMSR_JAX:007914) and Ai6(RCL-ZsGreen) (Jackson Laboratory, IMSR_JAX:007906) fluorescent indicator strains, respectively. The resulting two lines showed fluorescent labeling restricted to known cholinergic systems, including basal forebrain cholinergic neurons of the MS-diagonal band of Broca complex and large ChINs of the CPU.

### Microscopic assessment of fluorescent protein signals in formaldehyde-fixed sections

ChAT-tdTomato (N=3) and ChAT-ZsGreen (N=3) mice heterozygous for both the Cre and the indicator gene alleles were anesthetized with a cocktail of ketamine (25 mg/kg), xylavet (5 mg/kg), and pipolphen (2.5 mg/kg) in saline and perfused transcardially with 80 ml of ice-cold 0.5%, 2% or 4% formaldehyde (prepared freshly from paraformaldehyde) in 0.1 M phosphate buffered saline (PBS; 0.1 M; pH 7.4), followed by 50 ml of ice-cold 20% sucrose solution in PBS. The brains were removed, snap-frozen in powdered dry ice and stored at -80 °C until sectioned at 12 µm in the coronal plane with a Leica CM1860 UV cryostat (Leica Biosystems, Wetzlar, Germany). Sections corresponding to Atlas plates 20-26 of Paxinos (Bregma +1.34 to +0.62 mm) (35) were thaw-mounted on glass slides. Fluorescent images of CPU cholinergic neurons were captured with a Zeiss AxioImager M1 microscope (Carl Zeiss, Göttingen, Germany) using a 40× objective lens, filter sets for fluorescein isothiocyanate or rhodamine, an AxioCam MRc5 digital camera (Carl Zeiss) and the AxioVision Se64 Rel.4.9.1 software. The digital photomicrographs were prepared using identical manual exposure settings for all ChAT-tdTomato and ChAT-ZsGreen specimens fixed with varying formaldehyde concentrations. Editing with the Adobe Photoshop CS5 software (Adobe System Inc., California, USA) was applied to the composite table to maintain technical variations between samples.

### Transcriptomic studies

#### Section preparation for LCM

For all experiments, reagents were of molecular biology grade. Buffers were pretreated overnight with diethylpyrocarbonate (DEPC; Merck; 1 ml/L) and autoclaved or prepared using DEPC-treated and autoclaved water as diluent. The working area was cleaned with RNaseZAP (Merck KGaA, Darmstadt, Germany). To minimize possible adverse effects of fixation on RNA integrity and recovery, 0.5% formaldehyde in 0.1 M PBS (80 ml) was chosen for transcardiac perfusion of ChAT-ZsGreen mice, followed by ice-cold 20% sucrose in PBS (50 ml). The brains were snap-frozen on pulverized dry ice and stored at -80 °C until sectioned with a cryostat. Twelve-µm-thick sections containing the MS and the CPU (Atlas plates 20-26 of Paxinos; Bregma +1.34 to +0.62 mm) (35) were thaw-mounted onto PEN membrane glass slides (Membrane Slide 1.0 PEN, Carl Zeiss), air-dried in the cryostat chamber and preprocessed for LCM as reported elsewhere (18, 22, 23). In brief, the slides were immersed sequentially in 50% EtOH (20 sec), n-buthanol:EtOH (25:1; 90 sec) and xylene substitute:n-butanol (25:1; 60 sec). Then, they were stored at -80 °C in slide mailers with silica gel desiccants or processed immediately for LCM.

#### Laser capture microdissection

ChAT-ZsGreen neurons were microdissected individually and pressure-catapulted into 0.5 ml tube caps (Adhesive Cap 200, Carl Zeiss) with a single laser pulse using a 40× objective lens and the PALM Microbeam system and RoboSoftware (Carl Zeiss). Neurons pooled from the MS or the dorsal CPU were stored in the LCM tube caps at -80 °C until RNA extraction.

#### RNA extraction

RNA samples were prepared with the Arcturus Paradise PLUS FFPE RNA Isolation Kit (Applied Biosystems, Waltham, MA, USA).

#### RNA analytics

RNA yield and integrity number (RIN) were determined with the Agilent 2100 Bioanalyzer system using the Eukaryotic Total RNA Pico Chips and the 2100 Expert software (Agilent, Santa Clara, CA, USA). To obtain enough RNA for this test, each sample (N=3) included 500 cholinergic neurons isolated from the CPU. The calculated RNA yield/neuron and RIN values were provided as mean±SEM of the three samples.

#### RNA-Seq library preparation

Sequencing libraries were prepared with the TruSeq Stranded Total RNA Library Preparation Gold kit (Illumina, San Diego, CA, USA) from 300 cholinergic neurons microdissected and pooled from the dorsal CPU (CPU300; N=4) and, to achieve similar microdissected section areas, from 330 cholinergic neurons of the MS (MS330; N=4). Additional libraries were generated from 30 CPU cholinergic neurons (CPU30; N=4). The 15 PCR cycles recommended by the manufacturer for DNA fragment enrichment was increased to 16 using 300-330 neurons (18) and to 19 using 30 cells. A 1 nM library mix (20 µl) containing the twelve indexed samples proportionally was sequenced with an Illumina NextSeq500 instrument using the NextSeq500/550 High Output v2.5 kit (75 cycles).

#### Bioinformatics

Trimmomatic 0.38 (settings: LEADING:3, TRAILING:3, SLIDINGWINDOW:4:15, MINLEN:36) (36) and Cutadapt (37) were used to remove low quality and adapter sequences, respectively. Remaining reads were mapped to the Ensembl mm107 mouse reference genome using STAR (v 2.7.9a) (38). Read assignment to genes, read summarization and gene level quantification were performed by featureCounts (Subread v 2.0.2) (39). The counts per million read (CPM) values were calculated with the edgeR R-package (40). The coefficient of variation (CV) of CPM values for each of the four CPU300, CPU30 and MS330 transcriptomes were determined for the most abundant 1,000 transcripts and illustrated in violin plot. Transcriptome coverage was defined as the number of different transcripts (cutoff: reads≥ 5) in each sample and presented as the mean±SEM of four transcriptomes for each library type. RSeQC analysis (41) was used to analyze read distribution over gene body and the distribution of reads between exons, introns and the genomic space between annotated genes.

#### Species similarities in the receptor profile of striatal ChINs from the mouse dorsal CPU and the human putamen

Transcripts with the highest mean CPMs were identified in the CPU300 transcriptomes as well as in the analogous ChIN database of the human putamen we reported recently (17). The top 100 receptors (KEGG BRITE database of the Kyoto Encyclopedia of Genes and Genomes; KEGG) (42) in each species and the corresponding receptors in the other one, were illustrated with dot plots embedded in a Venn diagram.

#### Functional classification of MS330 and CPU300 transcripts

Transcripts present at CPM>1 in the CPU300 and MS330 transcriptomes were arranged by mean CPMs and ranked according to their presence in 4, 3, 2 or only 1 of the four samples. Selected functional categories (*ion channels, transporters, transcription factors* and *receptors*) were defined using the KEGG BRITE database.

#### Identification of neurodegenerative disease genes in cholinergic neurons

Identification of Alzheimer’s disease and Parkinson’s disease genes in the two spatially defined cholinergic neuron populations was based on the Human Disease Ontology database (43).

#### Differential expression analysis of cholinergic neuron populations

Differential expression analysis of the CPU and MS cholinergic cell types was performed via comparing the MS330 and CPU300 transcriptomes with the DESeq2 R-package (44), following correction for unwanted variations with RUVSeq (45). Differences in mRNA expression levels were quantified by log_2_ fold changes (log_2_FC), using approximate posterior estimation for GLM coefficients (Apeglm method) (46). To take multiple testing into account, p values were corrected by the method of Benjamini and Hochberg (47). Volcano plot and heat map, made with the EnhancedVolcano and Pheatmap program packages, respectively, were used to illustrate all transcripts that were expressed differentially in the spatially defined CPU300 and MS330 samples. Top 10 transcripts in the *ion channels, transporters, transcription factors* and *receptors* categories were illustrated in box plots. Synaptic gene ontology (SynGO) (48) analysis was used to classify synapse-related genes.

#### Studying the effects of limited RNA input of sequencing quality

Quality of sequencing in three types of transcriptome (MS330, CPU300 and CPU30), each including 4 independent samples) was determined from several technical parameters. These included total reads, aligned reads and assigned reads. Library homogeneity was assessed from the mean CV of the top 1,000 most abundant transcripts and from the results of principal component analysis (PCA).

In addition, homogeneity of heat maps was assessed for the top 1,000 CPU transcripts (highest mean CPMs in the 8 CPU transcriptomes). Transcriptome coverage was defined as the number of different transcripts in each library.

### Immunofluorescence experiments

#### Perfusion-fixation and section preparation

To provide morphological support for the roles of dopamine and TRH in the regulation MS and CPU cholinergic neurons, respectively, ChAT-ZsGreen mice (N=5) were anesthetized and sacrificed by transcardiac perfusion with 40 ml 4% formaldehyde in 0.1 M PBS (pH 7.4). The brains were removed, postfixed for 1h, infiltrated with 20% sucrose overnight, and then, snap-frozen on dry ice.

#### Section preparation

Twenty-μm-thick coronal sections containing the MS and the CPU were prepared with a Leica SM 2000R freezing microtome (Leica Microsystems) and stored at -20 °C in 24-well tissue culture plates containing antifreeze solution [30% ethylene glycol, 25% glycerol, 0.05 M phosphate buffer (pH 7.4)].

#### Fluorescent visualization of cholinergic neurons and their afferents

Floating sections were pretreated with 1% H_2_O_2_ and 0.5% Triton X-100 and then, processed for the immunofluorescent detection of either the dopaminergic marker enzyme tyrosine hydroxylase (TH) or proTRH. Cholinergic cells were recognized based on their ZsGreen fluorescence. Dopaminergic neuronal elements were detected with chicken anti-TH antibodies (AB_10013440, Aves Laboratories, Davis, CA, USA; 1:1,000; 48h; 4 °C) (49) and TRH neurons with rabbit proTRH antibodies (PA5-57331, Invitrogen, Waltham, MA, USA; 1:1,000; 48h; 4 °C) (50). Primary antibodies were reacted with Cy3-conjugated secondary antibodies (Jackson ImmunoResearch; 1:500; 2h; RT). The labeled sections were mounted on microscope slides, coverslipped with the aqueous mounting medium Mowiol and analyzed with confocal microscopy.

#### Confocal microscopy

Fluorescent signals were studied with a Zeiss LSM780 confocal microscope. High-resolution images were captured using 20×/0.8 NA and 63×/1.4 NA objective lenses, a 0.6–1× optical zoom and the Zen software (CarlZeiss). ZsGreen was detected with the 488 nm and Cy3 with the 561 nm laser line. Emission filters were 493–556 nm for ZsGreen and 570–624 nm for Cy3. Emission crosstalk between the fluorophores was prevented using ‘smart setup’ function. To illustrate the results, confocal Z-stacks (Z-steps: 0.85-1 μm, pixel dwell time: 0.79-1.58 μs, resolution: 1,024×1,024 pixels, pinhole size: set at 1 Airy unit) were merged using maximum intensity Z-projection (ImageJ). The final figures were adjusted in Adobe Photoshop using the magenta-green color combination and saved as TIF files.

### Slice electrophysiology

#### Brain slice preparation

Brain slices were prepared as described earlier (51) with slight modifications. Briefly, 20-30 day old male ChAT-Cre/ZsGreen mice (N=13) were decapitated in deep isoflurane anesthesia. The brain was immersed in ice-cold low-Na cutting solution bubbled with carbogen (mixture of 95% O_2_ and 5% CO_2_). The cutting solution contained the following (in mM): saccharose 205, KCl 2.5, NaHCO_3_ 26, MgCl_2_ 5, NaH_2_PO_4_ 1.25, CaCl_2_ 1, glucose 10. Brain blocks including the MS and the dorsal CPU were dissected. 220-μm-thick coronal slices were prepared with a VT-1000S vibratome (Leica Microsystems) and transferred into oxygenated artificial cerebrospinal fluid (aCSF; 33 °C) containing (in mM): NaCl 130, KCl 3.5, NaHCO_3_ 26, MgSO_4_ 1.2, NaH_2_PO_4_ 1.25, CaCl_2_ 2.5, glucose 10. The solution was then allowed to equilibrate to room temperature for 1 hour.

#### Whole cell patch clamp experiments

Recordings were carried out in oxygenated aCSF at 33 °C using Axopatch-200B patch-clamp amplifier, Digidata-1322A data acquisition system, and pClamp 10.4 software (Molecular Devices Co., Silicon Valley, CA, USA). Neurons were visualized with a BX51WI IR-DIC microscope (Olympus Co., Tokyo, Japan). ChAT-ZsGreen neurons showing green fluorescence were identified by a brief illumination (CoolLED, pE-100, Andover, UK) at 470 nm using an epifluorescent filter set. The patch electrodes (OD=1.5 mm, thin wall; WPI, Worcester, MA, USA) were pulled with a Flaming-Brown P-97 puller (Sutter Instrument Co., Novato, CA, USA). Electrode resistance was 2–3 MΩ. The intracellular pipette solution contained (in mM): K-gluconate 130, KCl 10, NaCl 10, HEPES 10, MgCl_2_ 0.1, EGTA 1, Mg-ATP 4, Na-GTP 0.3, pH=7.2-7.3 with KOH. Osmolarity was adjusted to 300 mOsm with D-sorbitol. Pipette offset potential, series resistance (R_s_) and capacitance were compensated before recording. Only cells with low holding current (< ≈50 pA) and stable baseline were used. Input resistance (R_in_), R_s_, and membrane capacitance (C_m_) were also measured before each recording by using 5 mV hyperpolarizing pulses. To ensure consistent recording qualities, only cells with R_s_<20 MΩ, R_in_>300MΩ, and C_m_ >10 pF were accepted.

Spontaneous firing activity of ChAT-ZsGreen neurons were recorded in whole-cell current clamp mode. Measurements started with control recording (1 min). Then a single bolus of the dopamine D2 receptor agonist sumanirole (30 μM, Tocris No.2773 Abingdon, UK) (52) or TRH (10 μM, Thermo Fisher [Bachem] No.4002023.0025 MA, USA) (53) was pipetted into the aCSF-filled measurement chamber and recording continued for another 7 min. Each neuron served as its own control when treatment effects were evaluated.

#### Statistical analysis

Recordings were stored and analyzed off-line. Event detection was performed using the Clampfit module of the pClamp 10.4 software (Molecular Devices Co., Silicon Valley, CA, USA). Mean firing rates were calculated from the number of action potentials (APs) over the control and treatment periods, respectively. All the recordings were self-controlled in each neuron and the effects were expressed as relative percentages of the control periods. Treatment group data were expressed as mean ± SEM. Statistical significance was determined with two-tailed Student’s *t*-tests using the Prism 3.0 software (GraphPad Inc., CA). Differences were considered significant at *p<*0.05.

## Acknowledgements

This research received funding from the Translational Neuroscience National Laboratory Program, the National Research, Development and Innovation Office (K128317 and 138137 to E.H. and PD134837 to K.S) and the Eötvös Loránd Research Network (SA-104/2021) of Hungary. We thank Ethan Tyler, Lex Kravitz and Emmett Thompson for open access science art at SciDraw (accessible at doi.org/10.5281/zenodo.3925901 doi.org/10.5281/zenodo.3925971).

## Data availability

RNA sequencing files generated in the current study are available in BioProject with the accession number PRJNA901862 (Release date: 04.30.2023).

For reviewing purposes: https://dataview.ncbi.nlm.nih.gov/object/PRJNA901862?reviewer=g1425pvom749pgh9snvhcla2ii

## Code availability

Scripts will be available upon request at https://github.com/goczbalazs/PRJNA901862

## Contributions

**Conceptualization**, É.R., B.G., M.S., K.S., S.T., I.F., S.P., M.P., N.S., E.H.;

**Methodology**, É.R., B.G.,

M.S., K.S., S.T., I.F., S.P., M.P., N.S., E.H.;

**Investigation**, É.R., B.G., M.S., K.S., S.T., I.F., S.P., M.P.,

N.S., E.H.;

**Writing – editing**, É.R., B.G., K.S., E.H.;

## Funding acquisition and supervision

K.S., E.H.

## Ethics declaration

Animal experiments were carried out in accordance with the Institutional Ethical Codex, Hungarian Act of Animal Care and Experimentation (1998, XXVIII, section 243/1998) and the European Union guidelines (directive 2010/63/EU) and approved by the Institutional Animal Care and Use Committee. All efforts were made to minimize potential pain or suffering and to reduce the number of animals used.

## Supporting Information for

**Supplementary Figure 1:**
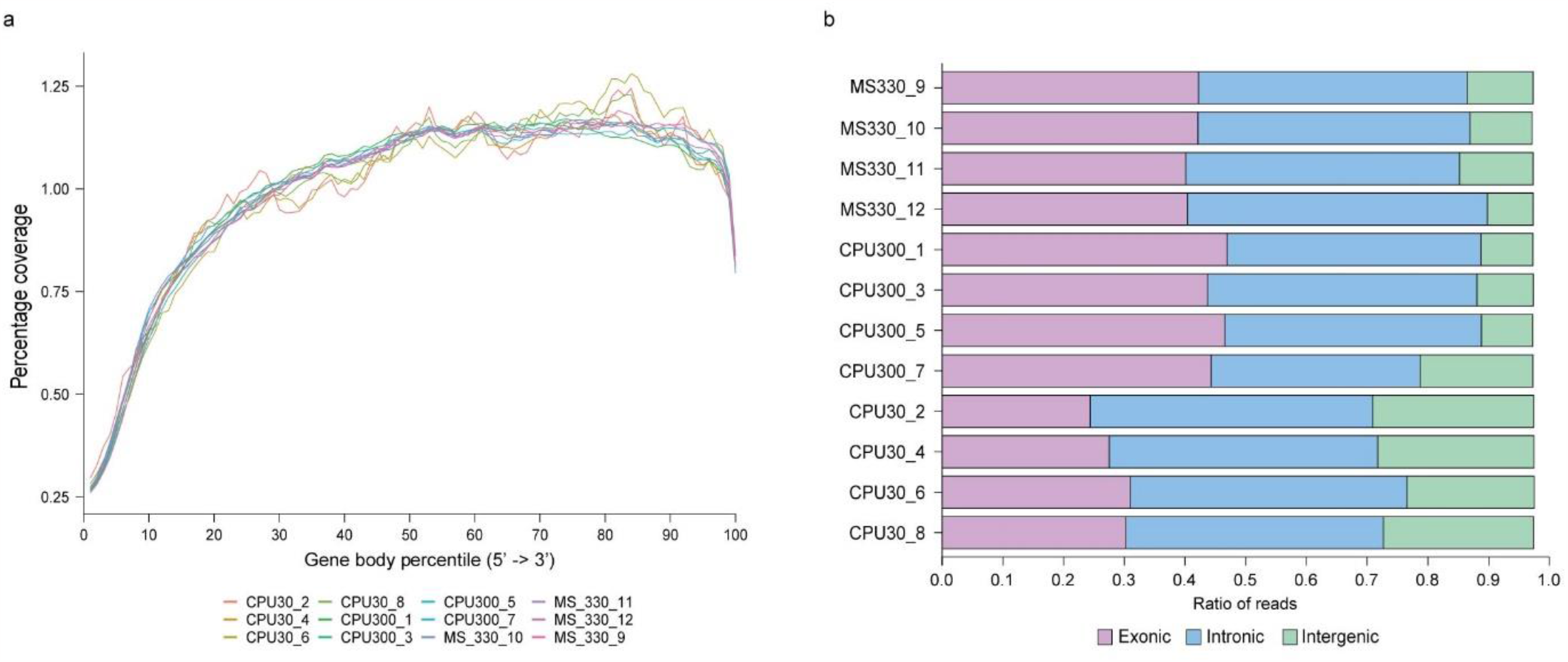
RSeQC analysis of reads. **a:** Read distribution over gene body plot. **b:** Distribution of transcript-associated reads that map within exons, introns and the genomic space between annotated genes. Note the reduced mapping of reads to exonic sequences and the increase in intergenic reads in the CPU30 samples.

**Supplementary Figure 2:**
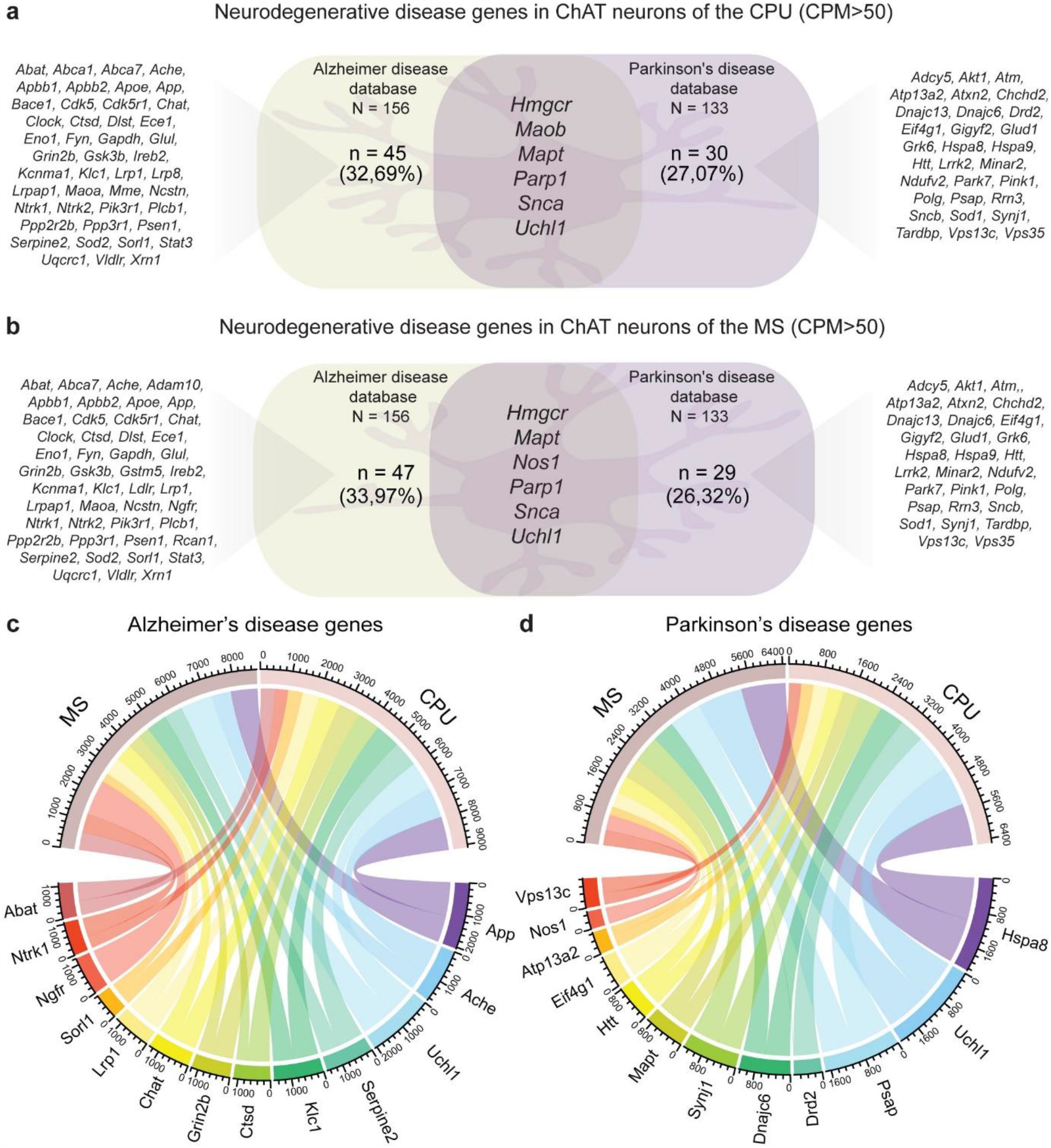
Enrichment of neurodegenerative disease genes in the two spatially defined cholinergic neuron populations. Cholinergic neurons both in the MS and the CPU highly express genes implicated in Alzheimer’s disease (AD) and Parkinson’s disease (PD). **a, b:** Venn diagrams reveal that ∼26-34% of genes listed in AD (N=156) and PD (N=133) Human Disease Ontology databases are expressed at high levels (CPM>50) in the two spatially distinct cholinergic neuron populations. **c, d:** Chord diagrams illustrate abundances (mean CPMs) of the top 10 most expressed AD genes (**c**) and PD genes (**d**) in each of the MS and CPU cholinergic cell types.

**Supplementary Figure 3:**
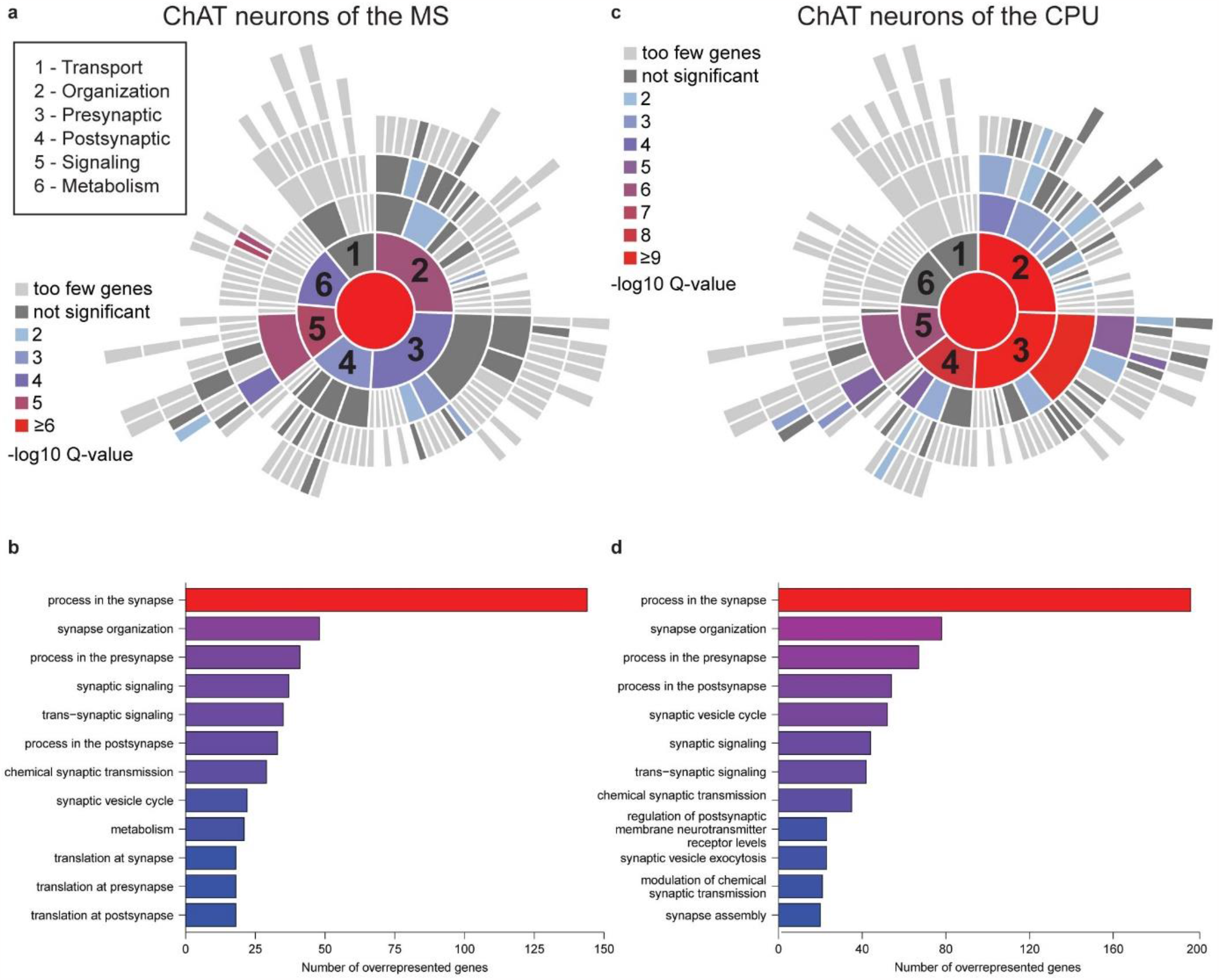
SynGO pathway analysis of differentially expressed genes. Many of the 2,891 transcripts expressed differentially between the MS330 and CPU300 transcriptomes are involved in synaptic functions. **a, b:** Synaptic genes expressed predominantly in MS cholinergic neurons. Diagrams graphically illustrate the number of genes in different Synaptic Gene Ontology (SynGO) categories. **c, d:** Synaptic genes with predominant expression in cholinergic neurons of the CPU. Specific genes are listed in **Source Data 2**.

**Supplementary Figure 4:**
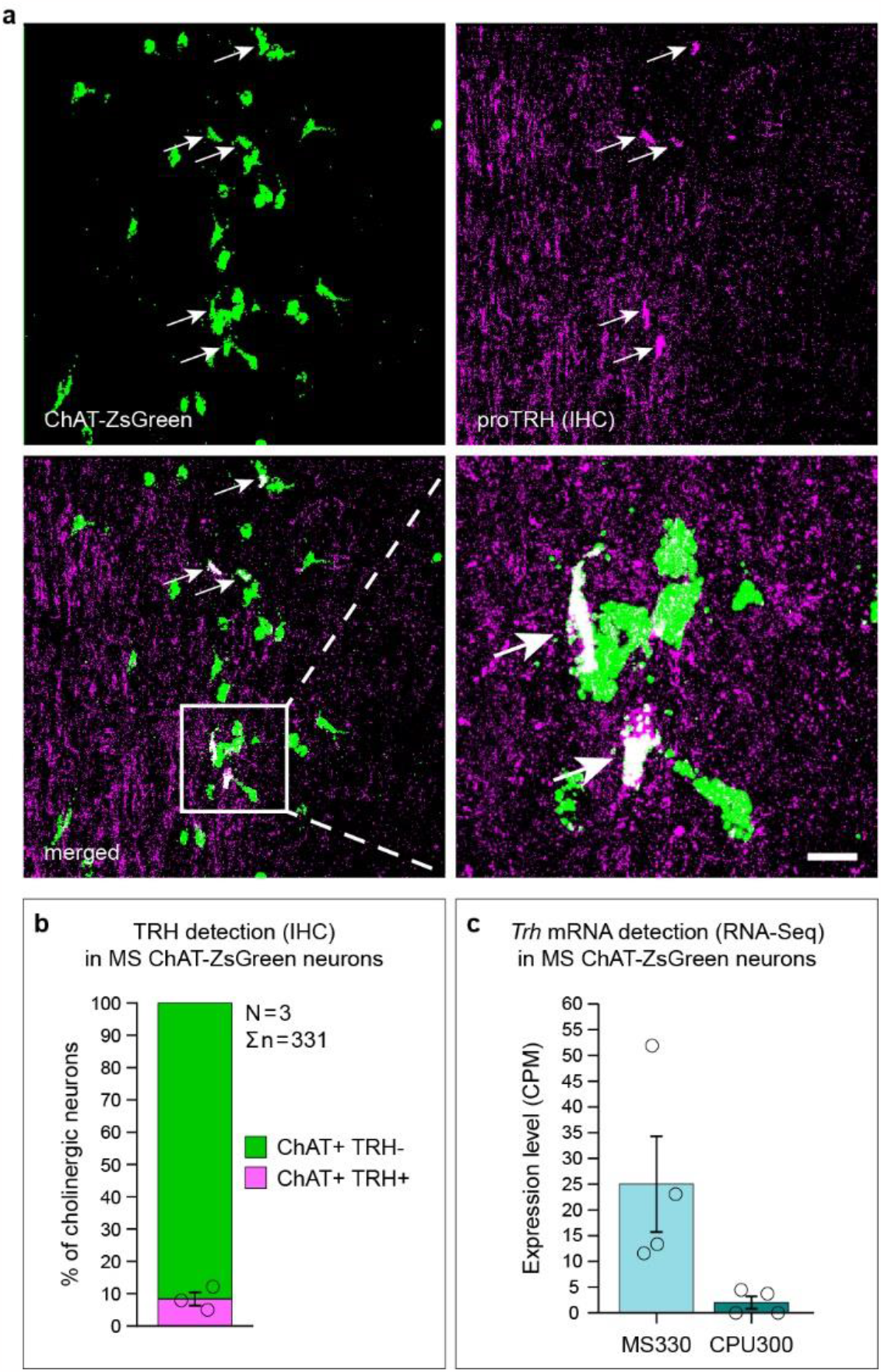
Results of anatomical and gene expression studies indicating TRH cotransmission in a subpopulation of MS cholinergic neurons. **a:** Confocal microscopic analysis of cholinergic (green; ChAT-ZsGreen signal) and TRH (magenta; proTRH immunoreactivity) neurons provides evidence that a subset of cholinergic neurons in the MS synthesize proTRH. Arrows point to double-labeled cell bodies. **b:** Quantitative analysis detects the proTRH signal in 8.4±2.1% (mean ± SEM) of MS cholinergic neurons. **c:** Presence of *Trh* mRNA in the CPU330 transcriptomes confirms that cholinergic neurons of the MS synthesize *bona fide* proTRH. Scale bar: 50 µm (**a**), 12.5 µm (high power inset in **a**).

